# Baseline activity of V1 interneurons connects pupil-linked arousal to engaged behavioral state

**DOI:** 10.64898/2026.02.05.704022

**Authors:** S.A. Nuiten, M.N. Oude Lohuis, A.C. Schaub, S. van Gaal, U. Olcese, C.M.A. Pennartz, P. Sterzer, J.W. de Gee

**Affiliations:** University Psychiatric Clinics, University of Basel, Basel, CH; Department of Clinical Research, University of Basel, Basel, CH; Champalimaud Neuroscience Programme, Champalimaud Foundation, Lisbon, POR; Department of Psychology, University of Amsterdam, Amsterdam, NLD; Amsterdam Brain & Cognition, University of Amsterdam, Amsterdam, NLD; Swammerdam Institute for Life Sciences, University of Amsterdam, Amsterdam, NLD

## Abstract

Humans and rodents alternate between discrete, persistent behavioral states during perceptual decision-making. For example, trials can be clustered into states of engaged, disengaged, and biased decision-making strategies. In mice, the probability of being in an engaged state exhibits an inverted-U relationship with baseline pupil-linked arousal, consistent with the Yerkes-Dodson Law. We replicated this relationship in mice (N=11; audio-visual change detection task) and showed that it generalizes to humans (N=69; auditory detection task). We then examined its neural basis, using simultaneous pupillometry and electrophysiological recordings in mouse V1. A nonlinear mediation model revealed that baseline activity of putative fast-spiking interneurons, but not putative pyramidal neurons, made a significant statistical contribution to the inverted-U shaped relationship between pupil-linked arousal and engaged-state probability. These findings are consistent with the idea that arousal may shape behavioral states by modulating baseline inhibitory activity within a primary sensory region.

## Introduction

We are all familiar with the contrast between the focused immersion of a “flow state”^1^ and periods of mind-wandering or distraction. Indeed, in challenging perceptual decision-making contexts, humans and mice alternate between discrete behavioral states with distinct choice behavior^2–8^. Example behavioral states, as revealed by generalized linear model-hidden Markov models (GLM-HMM), include “engaged” (choice behavior in line with external sensory stimuli), “biased” (choice behavior reflecting internal preferences), and “disengaged” (lower response rate and weak sensitivity to external stimuli). Because behavioral states profoundly impact how sensory inputs are translated to behavior, abnormal state dynamics may contribute to disorders characterized by epochs of distorted perception, such as schizophrenia^9–11^. This underscores the need to understand their physiological underpinnings.

The probability of being in an engaged state may be regulated by neuromodulatory nuclei in the brainstem, midbrain, and basal forebrain, whichtogether form the core of the ascending arousal system. They project to large parts of the cortex and change the functional properties of their target networks^12–17^. Consequently, neuromodulatory nuclei are ideally positioned to shape both local circuit dynamics and cortex-wide network activity underlying decision-making in a coordinated fashion. Pupil responses reflect the activity of multiple neuromodulatory systems^18–20^ and the ensuing cortical arousal state^21^. Critically, perceptual sensitivity and engaged-state probability peak during intermediate levels of baseline (pre-trial) pupil-linked arousal, and are reduced during low and high arousal states^4,5,22–27^, reminiscent of the classic Yerkes-Dodson law^28^.

Here, we examined whether baseline activity in primary visual cortex (V1) statistically mediates the relationship between arousal and choice behavior. Baseline V1 activity is a strong candidate for four reasons (but see ^6^): (i) V1 receives neuromodulatory input, especially from the noradrenergic locus coeruleus and cholinergic basal forebrain^29,30^, (ii) neuromodulatory input to V1 is known to shape neural response properties relevant for visual processing^31,32^, (iii) baseline V1 activity is minimal at intermediate levels of baseline pupil-linked arousal^33^, and (iv) baseline V1 activity is inversely related to task engagement^34–36^. Taken together, this leads to the hypothesis that baseline activity in V1 may account for the nonlinear relationship between pupil-linked arousal and engaged-state.

To directly test this hypothesis, we applied GLM-HMMs to session-wise time courses of choice behavior of mice and humans performing challenging perceptual decision-making tasks^37,38^. We replicated the previously observed “inverted-U” relationship between baseline pupil-linked arousal and engaged-state probability in mice^4,5^, demonstrated that this relationship generalizes to humans, and revealed that baseline activity of putative interneurons in mouse V1 statistically accounts for part of this nonlinear pattern.

## Results

### Pupil-linked arousal nonlinearly predicts engaged behavioral state in mice

We reanalyzed data of mice performing an audio-visual change detection task, in which they were continuously presented with an auditory stimulus (i.e., harmonic tone combinations) and a visual stimulus (i.e., moving grating) and were trained to report changes in tone frequency or grating orientation by licking one of two response spouts (**Figure 1A**)^37,38^. Auditory and visual changes occurred pseudo-randomly and at multiple saliency levels. We analyzed all behavioral data of expert mice for which pupillometry data was available on at least one session (N=11 mice, 143 sessions, 61029 trials). Across all trials, we observed robust psychometric functions indicating that correct response rates increased with the saliency of sensory changes (**Figure 1B**), showing that mice appropriately responded to task-relevant features of the cross-modal continuous stimulus.

**Figure 1.**
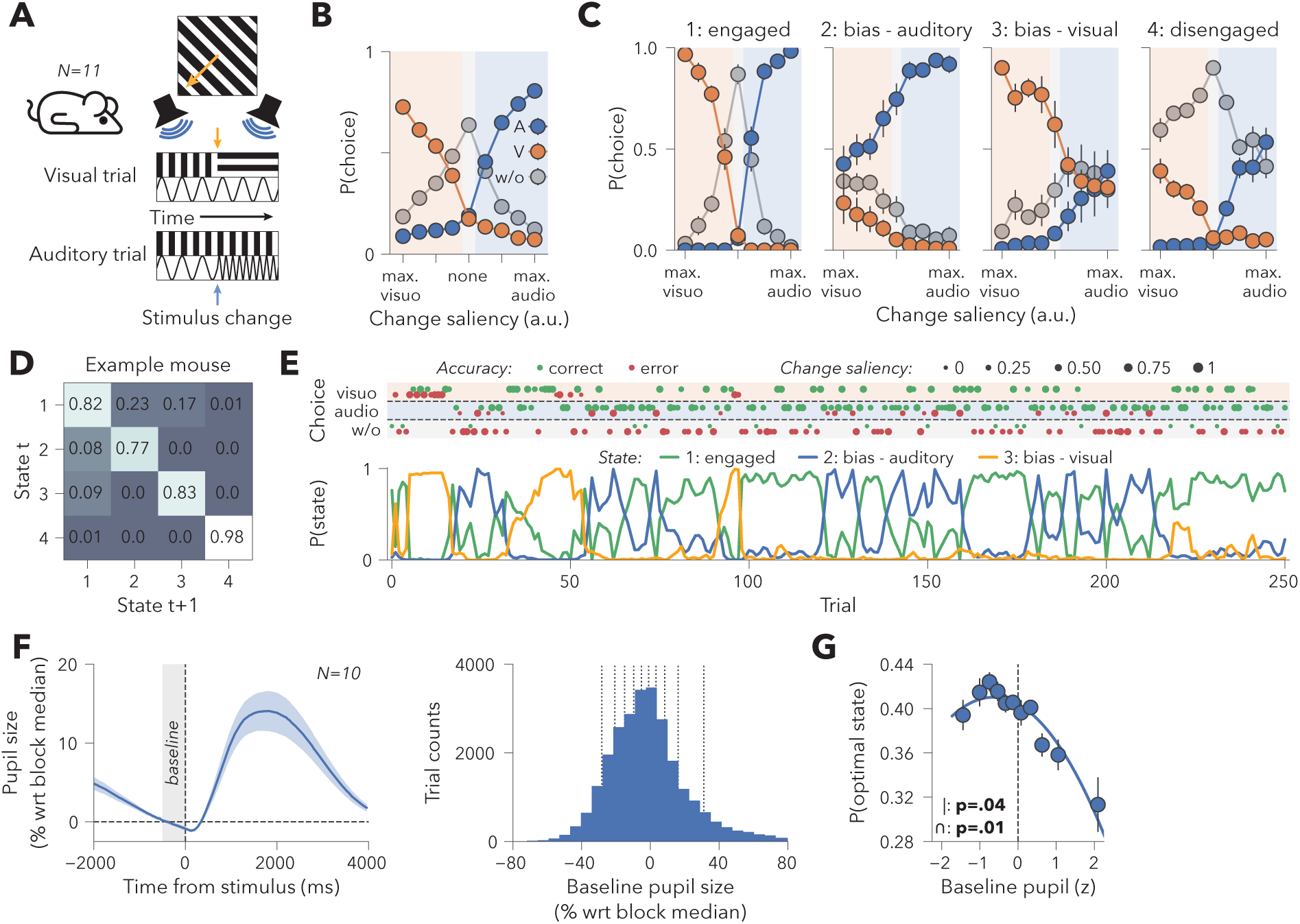
Engaged behavioral state probability is nonlinearly related to baseline pupil size in mice. **(A)** Schematic of audio-visual change detection task performed by mice (N=11 mice). **(B)** Average response rates across animals and all behavioral trials, separated by auditory responses (blue; “A”), visual responses (orange; “V”), and no-response trials (grey; “w/o”). Error bars are shown but too small to be visible. **(C)** Average response rates across animals for each of the four latent behavioral states. **(D)** Transition matrix describing transition probabilities for example mouse. High values on the diagonal indicate state persistence. **(E)** State dynamics during example session of the same mouse of panel D, showing transitions between persistent and discrete decision-making strategies. **(F-G)** Pupillometry analyses (N=10 mice; see main text). **(F)** *Left panel*: Average pupil response around the time of a stimulus change. Grey window indicates the period used to calculate baseline pupil size. *Right panel*: Histogram of baseline pupil sizes. Vertical dashed lines indicate the thresholds for pupil bins reported in panel G. Baseline pupil sizes were positively skewed, with more relatively large pupil sizes than relatively small pupil sizes (t(9)=5.05, p<.001). **(G)** Relationship between baseline pupil size and engaged-state probability. Linear mixed-effects model fitting was performed on single-trial z-scored data (Methods), pupil binning was only used for visualization. Individual-level polynomial regressions revealed significant quadratic relations between baseline pupil size and engaged-state probability for 9 out of 10 mice. Error bars and shading indicate SEM across animals.

We verified that mice performing the audio-visual change detection task exhibited persistent behavioral states. We used a GLM-HMM to discover sudden changes in decision strategies^2–5^. The GLM component modelled choices using an intercept (capturing choice bias) and stimulus input predictor (capturing sensitivity) and the HMM component described the single-trial transition probabilities between a fixed number of states that differed with respect to their GLM beta weights (**Methods**). The best fitting GLM-HMM (selected using cross-validation; **Figure S1A**) contained four HMM-states, including an engaged state with steep psychometric functions for both visual and auditory changes, two states in which responses were biased towards either auditory or visual choices, and a disengaged state with frequent behavioral lapses (**Figure 1C**). Behavioral states were persistent: state repetition probabilities were high (**Figure 1D**), resulting in states that often lasted several trials before transitioning (mean ± SD state duration= 43.80 s, SD=18.01 s; **Figure 1E**). Task disengagement was accompanied by a relative reduction in spontaneous licking prior to stimulus change onset, whereas biased states saw a relative increase in spontaneous licking behavior (**Figure S1**). State descriptives, unclassified states, individual-level states, and transition matrices are reported in **Figure S1B-D**. In sum, in line with earlier reports^2–6^, mice cycled through discrete and persistent behavioral states that were characterized by different decision strategies.

We next sought to replicate earlier work showing that engaged-state probability and baseline pupil size are non-monotonically related^4,5^. We excluded 58 sessions without available pupillometry recordings and 11 sessions in which engaged-state probability fell outside the range of 0.05 to 0.95 (final sample: N=10 mice, 74 sessions, 26111 trials). Baseline pupil size was calculated as the average pupil size in the 500 ms preceding stimulus (change) onset (**Figure 1F**, grey window; **Methods**)^22,25,39,40^.

To characterize the shape of the relationship between baseline pupil size and engaged-state probability, we compared two mixed-effects models (**Methods**) in which engaged-state probability was predicted by pupil size either linearly or also with an added quadratic term. The quadratic model was strongly favored (ΔAkaike Information Criterion [AIC]=-13.46). This model confirmed that engaged-state probability was predicted by baseline pupil size following an inverted-U relationship (**Figure 1G**). The decline in engage-state probability for very small pupils was less pronounced, but in line with earlier work^23,27^, possibly reflecting the limited number of trials at this end of the pupil range. Indeed, the distribution of trial-wise baseline pupil size was positively skewed across mice, indicating that relatively large pupil size measurements were more frequent than relatively small ones (**Figure 1F**). Nonetheless, in our data too, the probability of being in an engaged state was nonlinearly related to pupil-linked arousal.

### Pupil-linked arousal also nonlinearly predicts engaged behavioral state in humans

Next, we characterized whether the nonlinear relationship between pupil-linked arousal and engaged-state probability generalizes to humans. Perceptual sensitivity of human observers peaks during intermediate pupil-linked arousal^22,25^, but it is unknown if this is due to increased engaged-state probability. For example, healthy human participants may not exhibit many state transitions in typical experimental blocks. Human participants (N=69, one session per participant, 55219 trials) performed a yes/no auditory detection task, in which, on each trial, they reported whether they had heard a pure tone (300 Hz) hidden in dynamic auditory noise (**Figure 2A**). Across all trials, humans performed this task with high accuracy (**Figure 2B**).

**Figure 2.**
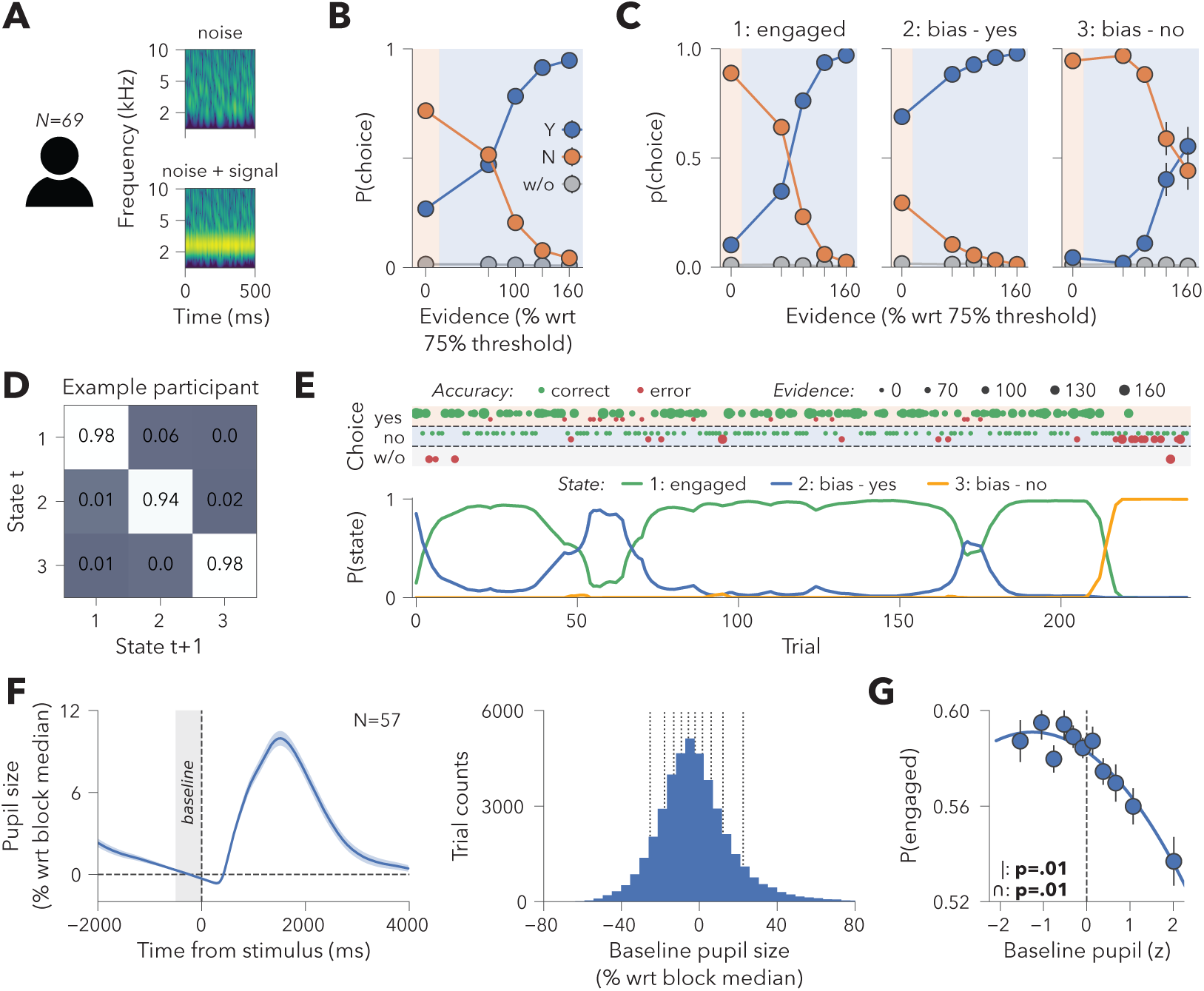
Nonlinear relationship between baseline pupil size and engaged-state probability generalizes to humans. **(A)** Schematic of auditory yes/no detection task performed by human participants (N=69 humans). **(B)** Average response rates across participants and all behavioral trials, separated by “yes” responses (blue; “Y”), “no” responses (orange; “N”), and no-response trials (grey; “w/o”). Error bars are shown but too small to be visible. **(C)** Average response rates across participants for each of the three latent behavioral states. Note that humans did not reveal a disengaged state, as they almost never had behavioral lapses. **(D)** Transition matrix describing for an example participant. High values on the diagonal indicate state persistence. **(E)** State dynamics during example session of the same participant of D, showing transitions between persistent and discrete decision-making strategies. **(F-G)** Pupillometry analyses (N=57 humans). **(F)** *Left panel*: Average pupil response around the time of a stimulus change. Grey window indicates the period used to calculate baseline pupil size. *Right panel*: Histogram of baseline pupil sizes. Vertical dashed lines indicate pupil binning thresholds. Vertical dashed lines indicate the thresholds for pupil bins reported in panel G. Baseline pupil sizes were positively skewed, with more relatively large pupil sizes than relatively small pupil sizes (t(56)=13.93, p<.001). **(G)** Relationship between baseline pupil size and engaged-state probability. Model fitting was performed on continuous z-scored data, pupil binning was only used for visualization. Individual-level polynomial regressions revealed significant quadratic relations between baseline pupil size and engaged-state probability for 34 out of 57 participants. Error bars and shading indicate SEM across participants.

We repeated the same analyses as for the mouse dataset. For humans, the best fitting GLM-HMM (selected using cross-validation; **Figure S2A**) contained three HMM-states, including an engaged state and two states in which responses were biased towards either yes or no choices (**Figure 2C**). There was no evidence of a disengaged state, consistent with the virtual absence of behavioral lapses, indicating that participants complied with the task-instruction to always provide a response (**Figure 2B**). Behavioral states were more persistent than in mice, suggesting humans were better able to maintain decision strategies over time (mean ± SD state duration=287.55s ± 303.69s; **Figure 2D-E**). State descriptives and unclassified states are reported in **Figure S2B-C**.

We excluded 12 human participants from subsequent analyses because their average engaged-state probability fell outside the range of 0.05 to 0.95 (final sample: N=57 human participants, 41235 trials). Engaged-state probability was best predicted by a mixed-level model including both linear and quadratic pupil predictors (ΔAIC=-38.89) and indeed displayed an inverted-U relationship with baseline pupil size (**Figure 2F-G**). Similar to the mice, the left limb of the curve declined less than the right, possibly reflecting the relatively small number of trials with relatively small pupil sizes due to the positive skew of baseline pupil sizes (**Figure 2F**). In sum, these findings show that humans also alternate between persistent behavioral states and that the nonlinear relationship between pupil-linked arousal and engaged-state probability is preserved across species.

### Baseline activity of fast-spiking interneurons in V1 statistically mediates the nonlinear relationship between pupil-linked arousal and engaged-state probability

Finally, we tested our hypothesis that baseline spiking activity in mouse V1 statistically mediates the relationship between pupil-linked arousal and engaged-state probability. We analyzed all behavioral data of sessions that included simultaneous pupillometry and extracellular recordings in V1 and applied several further criteria to ensure high data quality (final dataset: N=7 mice, 17 sessions, 4698 trials; **Methods**). In total, we included 95 putative fast-spiking interneurons (range, 3-11 per session; from here on called interneurons) and 172 putative pyramidal neurons (range, 4-21 per session; from here on called pyramidal neurons) (**Figure S3A**; **Methods**). Several key patterns were observed in this reduced data set: (i) pupil-linked arousal non-monotonically predicted engaged-state probability (**Figure 3A**), (ii) both putative cell-types exhibited robust “early” responses evoked by visual changes (**Figure 3B**)^37^, and (iii) both putative cell-types exhibited robust “late” responses on visual hit versus miss trials (**Figure 3B**)^37^.

**Figure 3.**
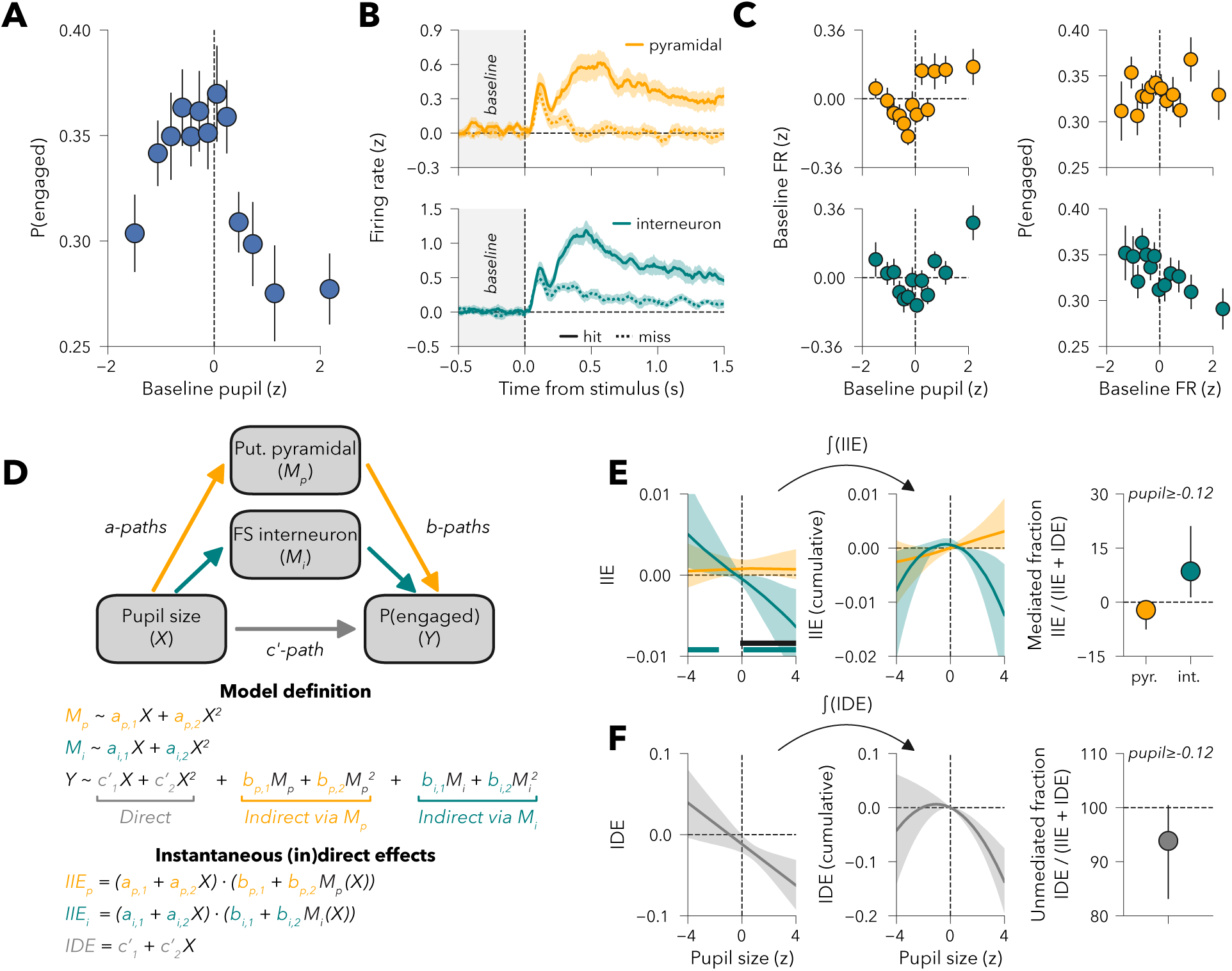
Nonlinear relationship between baseline pupil-linked arousal and engaged behavioral state is mediated by baseline V1 interneuron spiking activity. **(A)** Relationship between z-scored baseline pupil size and engaged-state probability (P(engaged)). **(B)** Normalized firing rate (FR; z-scored change from baseline) of V1 putative pyramidal neurons (*top*, orange) and GABAergic interneurons (*bottom*, teal) for hit and miss responses across all trials with a visual change. Grey window indicates the period used to calculate baseline firing rate. **(C)** Relationships across all trials between (*left*) baseline pupil size and z-scored baseline firing rates of V1 interneurons and putative pyramidal neurons and (*left*) z-scored baseline firing rates and P(engaged). **(D)** Schematic of the nonlinear mediation analysis. (E) *Left:* Instantaneous indirect effect (IIE) of pupil-linked arousal on P(engaged) via V1 spiking activity. Horizontal lines denote arousal levels where the IIE was significantly different from zero (teal, interneurons) or significantly differed between neuron types (black). *Middle:* Cumulative indirect effects, obtained by integrated the IIE over the full range of pupil sizes. *Right:* Proportion of the total arousal effect mediated by V1 spiking activity, averaged across arousal levels where the IIE differed significantly between neuron types. **(F)** Same as panel E, but for the instantaneous direct effect (IDE). Shading and error bars in panels A-C reflect SEM over sessions and in panels E-F reflect bootstrapped 95% confidence intervals.

We assessed the role of baseline V1 spiking activity separately for excitatory putative pyramidal cells and interneurons, because of (i) the different roles they play in information processing^41^, (ii) the different neuromodulatory receptor densities they express on their membranes^42^, (iii) the cell-type specific relationships between evoked activity and pupil-linked^43^, (iv) a recent model positing a specific role for interneurons in this context^22^, and (v) recent work that established that GABAergic interneurons in V1 control neural and perceptual contrast sensitivity^44^. We applied a statistical nonlinear mediation model, which isolates the portion of the arousal-engagement relationship uniquely associated with V1 activity, providing insight beyond what pairwise analyses can reveal. Importantly, this approach is correlational in nature and does not assume or impose any underlying causal structure.

To illustrate the qualitative structure of the data, we first visually inspected the relationships between baseline pupil size, baseline V1 spiking activity, and engaged-state probability (formal statistical tests are reported below and in **Figure S3D**). We calculated baseline activity as the mean firing rate during the 500 ms preceding stimulus change, using the same window as for pupil size. For each session, trials were divided into thirteen equally sized bins based on baseline pupil size and baseline spiking activity. In line with earlier work^33–36^, we observed a U-shape relationship between baseline pupil size and interneuron spiking activity, while engaged-state probability was highest on trials with the weakest baseline interneuron spiking activity (**Figure 3C** – bottom row). Increased baseline V1 interneuron activity during non-engaged states could not be explained by an increased lick rate, as licking behavior was relatively elevated during the engaged state compared to all other states (**Figure S3B**).

The three visually apparent relationships—the U-shape between baseline pupil size and interneuron spiking, the inverted-U between baseline pupil size and engaged-state probability, and the negative link between interneuron spiking and engaged-state probability—form a coherent pattern: pupil-linked arousal is associated with elevated baseline interneuron spiking at high and low levels, and high interneuron activity corresponds to lower engagement. This is the qualitative structure expected if arousal modulates behavioral state through its impact on V1 interneurons. For pyramidal neurons, these patterns were less clear (**Figure 3C** – top row).

To formally test whether the relationship between baseline pupil size (predictor; *X*) and engaged-state probability (output; *Y*) was mediated by changes in baseline V1 spiking activity (mediator; *M*), we constructed a nonlinear mediation model (**Figure 3D**). Each model path (*a*, *b*, *c’*) included linear and quadratic terms to capture potential nonlinear effects. In mediation, the indirect effect is calculated as the product of the derivative of the *a*-path (𝑋 → 𝑀) and the *b*-path (𝑀 → 𝑌). In the linear case, this is equivalent to simply multiplying the beta coefficients of the *a*-path and *b*-path. In the nonlinear case (as used here), both derivatives can be expressed with respect to *X* (**Figure 3D; Methods**). The resulting product, the indirect effect, calculated across the full dynamic range of X, has been termed the instantaneous indirect effect (IIE)^45^, which reflects how the magnitude and direction of mediation may vary across baseline pupil-linked arousal states (**Methods**). The corresponding instantaneous direct effect (IDE) quantifies the portion of the 𝑋 → 𝑌 relationship that is independent from mediation by V1 activity. The model was jointly fit with interneurons and pyramidal cells as separate mediators on single-trial data (see **Figure S3D** for fitted beta values).

The resulting IIE confirmed our hypothesis: on trials characterized by low baseline pupil size (z-scored pupil ≤ -1.71), increases in pupil-linked arousal were associated with higher engaged-state probability via interneuron firing (positive IIE), where this relationship was inverted on trials characterized by larger baseline pupil size (z-scored pupil ≥ 0.11; **Figure 3E** – left panel). Integrating the interneuron IIE across the full dynamic range of pupil size shows that V1 interneuron activity contributes to the inverted-U relationship between baseline pupil-linked arousal and engaged-state probability (**Figure 3E** – middle panel). In contrast, pyramidal neurons showed no significant IIE at any level of baseline pupil size (**Figure 3E** – left panel). The mediation effect via interneuron activity at intermediate-to-large pupil sizes (pupil ≥ -0.12) was also significantly greater than that via pyramidal neuron firing (**Figure 3E** – left panel).

Finally, we assessed the magnitude of the mediation effect for interneurons at these high arousal levels. We calculated the mediated proportion of the total effect, defined as the sum of both IIEs and the IDE (**Figure 3F**). Approximately 8.53% (bootstrapped 95% confidence interval, CI_95_=[1.36, 21.11]) of the total path was explained by the interneuron-mediated path, whereas -2.17% (CI_95_=[-7.57, 0.20]) was attributed to the putative pyramidal neuron path (**Figure 3E** – right panel). The negative proportion for pyramidal IIE may suggest that this pathway opposed the direction of the total effect, as can also be seen in the positive pyramidal IIE at high levels of arousal (**Figure 3E** – middle panel). However, because the pyramidal IIE was not significant, this pattern is best interpreted as sampling variance rather than evidence for a substantive suppressor effect.

Together, these findings are consistent with the idea that arousal may shape behavioral states by modulating baseline inhibitory activity in V1.

## Discussion

Mouse and human perceptual choice behavior is typically not stationary but characterized by alternations between persistent yet distinct behavioral states in which different decision strategies are employed^2–6^. We found that these state dynamics are systematically related to baseline pupil-linked arousal, with the probability of occupying an engaged behavioral state peaking at intermediate arousal levels in both species. Electrophysiological recordings in mouse V1 further revealed that baseline activity of putative interneurons, but not putative pyramidal neurons, nonlinearly mediated the arousal-engagement relationship. Together, these findings advance two major conclusions: (i) arousal serves as a conserved, cross-species influence on perceptual decision-making by shaping engaged-state probability and (ii) this influence is, in part, related to the tuning of local inhibitory circuits in V1.

Because the relationship between arousal and engaged state is conserved across humans and mice, an important question is what key evolutionary advantage this mechanism provides. One possibility is that remaining in an engaged state is energetically costly, and organisms use a cost-benefit analysis to regulate arousal level and the ensuing behavioral state^46^. The outcome of this cost-benefit analyses may vary as a function of fatigue, satiety, and other factors. Another, not mutually exclusive scenario, is that organisms strike a balance between exploitation and exploration^47,48^, where biased and/or disengaged behavioral states may map onto exploratory behaviors to try and maximize reward. In both cases, orbitofrontal cortex and dorsal anterior cingulate cortex are likely candidate effectors, as both perform value computations to optimize behavior. These structures are strongly connected to sensory cortices and to relevant neuromodulatory nuclei^12,20,49–51^.

Impaired regulation of arousal and its influence on behavioral state dynamics may contribute to neurodevelopmental disorders such as ADHD^52^ and autism^53^. In ADHD, for example, fluctuations in arousal-linked engagement could underlie periods of inattentiveness and distractibility. Altered arousal-state dynamics may also be relevant for psychiatric disorders characterized by episodes of distorted perception, including schizophrenia^54–56^. Schizophrenia patients and healthy individuals with schizotypal traits exhibit reduced transient pupil-linked arousal responses to decision uncertainty^57–59^, suggesting that the networks associated with uncertainty processing^60,61^ may be less able to modulate arousal for optimizing behavior. Future work should determine whether changes in arousal contribute to state-like effects on attention and perception in these conditions and whether targeted modulation of these systems can support more stable perceptual and cognitive function.

Although we identified contributions of baseline V1 interneuron activity to the arousal-engagement relationship, the arousal-dependent modulation of behavioral states likely extends to other brain regions involved in decision-making, including other primary sensory regions, higher-order sensory regions, or associative cortices. Indeed, there is evidence for an inverted-U relationship between pupil-linked arousal and stimulus-evoked neural activity in auditory cortex: at intermediate baseline pupil size (i) tone-evoked excitatory postsynaptic potentials and spike rate are highest and most reliable^23^, and (ii) the neural discriminability of tones is maximal^24^. Recent work further shows that arousal-modulations of decision strategy are also associated with activity measured over motor cortex, reflecting later stages of decision formation and response preparation^39^. In line with this, a recent report describes that activity in posterior parietal cortex, but not sensory cortex (auditory), is causally related to shaping behavioral states^6^. However, this task required mice to traverse through a T-maze by running on a spherical treadmill, so it is likely that mice were on average in a higher state of arousal and engagement compared to the stationary head-fixed. Therefore, it remains an open question whether the three-way association between baseline arousal, baseline V1 interneuron activity, and behavioral state extends to other cortical regions or reflects broader network-level dynamics.

While this study provides valuable insights into the link between baseline pupil-linked arousal, baseline V1 spiking activity and behavioral states, several limitations should be acknowledged. First, we used pupil responses as a peripheral readout of changes in arousal state^21,62^. Changes in pupil diameter have been associated with locus coeruleus (LC) activity in humans^20,63,64^, monkeys^62,65^, and mice^19,66,67^. However, some (of these) studies also found unique contributions to pupil size from other neuromodulatory structures, such as the cholinergic basal forebrain^19,20,64,68^, the dopaminergic midbrain^20,64^, and serotonergic raphe nuclei^64,67,69^. Thus, future studies should pinpoint the exact neuroanatomical and neurochemical source(s) of our observed effects. Second, the physiological specificity of our extracellular measurements was insufficient to discriminate between different interneuron subtypes. Arousal-dependent modulation of neural responses in auditory cortex is cell-type specific, with excitatory and inhibitory subtypes exhibiting varying arousal-dependent activity patterns^43^ and a recent neurobiologically plausible model posits that the relationship between arousal and behavior is mediated by a disinhibitory circuit composed of interneurons expressing somatostatin and vasoactive intestinal peptide^22^. Future experiments using opto-tagging or 2p-imaging should characterize the types of interneurons and their interactions relevant to this context. Third, our results are correlational, and it remains to be determined if baseline V1 interneuron spiking is causally driven by pupil-linked arousal and shapes behavioral states. Finally, though mice and humans both made immediate binary choices following sensory input, the specific behavioral tasks differed, limiting the extent to which findings can be compared across species. Together, these limitations highlight avenues for future studies to clarify the circuit- and system-level mechanisms linking arousal, cortical activity, and behavior across various species.

Our research highlights the importance of conducting comparative experiments between humans and non-human species. It is generally assumed that the fundamental functions of arousal systems are similar in both humans and rodents. However, the exact way these systems function during decision-making has remained uncertain. Recent findings have demonstrated that both rodents (rats) and humans accumulate perceptual evidence in a similar way^70^, and that this process is influenced by phasic pupil-linked arousal in a comparable manner^71^. Our findings suggest that the regulation of behavioral state by baseline levels of pupil-linked arousal follows a principle that is also conserved across species.

## Methods

### Subjects

#### Mouse dataset

We reanalyzed data from a previous study^37,38^. From the total sample of 49 mice, 11 mice were selected based on the presence of pupillometry recordings during at least one experimental session. The animal experiment was performed according to national and institutional regulations, and the experimental protocol was approved by the Dutch Commission for Animal Experiments and by the Animal Welfare Body of the University of Amsterdam. A detailed description of the transgenic mouse lines, animal treatment, and head-bar surgery can be found in the initial publication^37^.

#### Human dataset

For the human dataset, 69 healthy participants (50 female, 19 male) were recruited from the online research environment of the University of Basel. Participants were aged between 18-56 (μ=26.71, sd=8.81). Before inclusion, participants were screened for psychosis-proneness with the Peter’s Delusion Inventory^72^ and included in the study when their PDI-score fell below 3 (low psychosis-proneness) or above 11 (high psychosis-proneness). Psychosis-proneness was not included as a variable of interest in the current study. This study was approved by the Ethics Committee for Northwest and Central Switzerland (EKNZ). Written informed consent was obtained from all participants after explanation of the experimental protocol. All participants received a monetary compensation of 20 Swiss Francs per hour for participation in this study.

### Behavioral tasks

#### Mouse dataset – audio-visual change detection task

Mice were continuously presented auditory and visual stimuli during an experimental session. Visual stimuli consisted of full-field drifting square-wave gratings that were presented with at 60Hz on an 18.5-inch monitor at 21cm from the eyes of the mouse. Gratings were presented with a temporal frequency of 1.5Hz and spatial frequency of 0.08 cycles per degree at 70% contrast. Auditory stimuli were combinations of five pure tones at harmonic frequencies. For example, a stimulus could be made up out of a central frequency of 2^13^Hz, with surrounding harmonics at 2^11^Hz, 2^12^Hz, 2^14^Hz, and 2^15^Hz. There were four categories of trials: visual trials in which the orientation of the grating changed, auditory trials in which the frequency of the 5 pure tones changed, catch trials in which both visual and auditory stimuli did not change, and conflict trials in which both visual and auditory stimuli changed. Mice reported the presence of visual and auditory changes by licking one of two spouts located in front of them. After a visual or auditory change, the respective stimulus feature (grating orientation, tone frequency) remained unaltered until the next visual or auditory change. The degree of orientation change determined the visual change saliency and the degree of change in tone frequency/octave determined the auditory change saliency.

Mice were subjected to a water restriction schedule and could earn their daily ration of liquid by performing the behavioral task. Food was readily available without limitations for the mice outside experimental sessions. Mice were head-fixed in a custom-build head-bar holder within a darkened and sound-isolated cabinet. Bodily movements were minimized by placing the mouse body in a small tube. Details on the behavioral training procedure are explicated in the original publication^37^. In total, 11 mice completed 143 sessions (8-21 sessions per mouse) and 61029 trials (160-799 trials per session).

#### Human dataset - Auditory detection task

Human participants performed an auditory yes/no detection task, in which they had to detect a pure tone (300Hz) that was presented for 500ms on top of auditory noise (temporally orthogonal ripple combination stimulus; TORC)^73^ on 50% of all trials. “No” decisions were given by pressing the “A” or “S” keys on a keyboard with the left hand, “yes” decisions were given by pressing the “K” or “L” keys with the right hand. Simultaneous with their first-order perceptual decision, participants indicated the binary confidence in their decision. For both hands high confidence was indicated with the middle finger and low confidence with the index finger. For example, high confidence “yes” responses were given by pressing “L” with the right middle finger. Confidence ratings were not analyzed in the current study. While participants performed the yes/no detection task, they were instructed to fixate on a fixation mark (black dot 0.26°) that was centrally presented on a gray background.

Each trial consisted of a baseline interval (1000ms), followed by the stimulus interval (500ms), a response interval (maximally 2000ms, aborted after the response), a feedback interval (1000ms), and a variable inter-trial-interval of 1500-3500ms. Participants received visual feedback whenever they failed to respond in time (red fixation mark). Target stimuli were presented at varying volumes, based on the subject-specific task difficulty (see below). Specifically, targets were presented at 70% (on 15% of all trials), 100% (on 15% of all trials), 130% (on 15% of all trials), and 160% (on 5% of all trials) of the titrated volume. The noise stimulus was presented in isolation on the remaining 50% of all trials. Participants performed minimally 720 trials of this task divided over three larger blocks, each consisting of mini-blocks of 60 trials that lasted approximately 5 minutes. Breaks between mini-blocks were self-paced.

Participants performed the experiment at the research facility of the University Psychiatric Clinics Basel. They were seated in a darkened, sound isolated room, resting their chins on a head-mount located 60 cm from a 53×30 cm screen (frequency: 100 Hz, resolution: 1920×1080). The main task and staircase procedure were programmed in Python 3.8 using PsychoPy^74^ and in-house scripts.

Before the primary task, participants completed a staircasing procedure (weighted up-down method^75^) that titrated task difficulty to 75% choice accuracy and included trial-by-trial feedback that helped participants familiarize themselves with the task.

##### Eye data acquisition in mice and humans

In mice, pupil size of the left eye was recorded at 25Hz using a near-infrared monochrome camera placed approximately 30cm from the mouse. Pupil recordings were available for 11 mice, 85 sessions (1-13 sessions per mouse), and 29957 trials (145-643 trials per session).

In humans, pupil diameter was continuously measured throughout the experiment at 1000Hz with an EyeLink 1000 eye tracker with an average spatial resolution of 15 to 30 min arc, using an EyeLink 1000 Long Range Mount (SR Research, Osgoode, Ontario, Canada). Trials were marked as faulty when the pupil signal was lost due to blinks or when participants’ gaze diverted ±1.5° from the fixation mark during the baseline interval, stimulus interval or response interval. Participants were presented with visual feedback whenever the pupil signal was lost (white fixation mark). Faulty trials were replaced with new trials, that were appended to the end of each block. This procedure ensured that each participant completed 720 useable trials. The eye tracker was calibrated before each block and participants were instructed not to move or reposition their heads after calibration until the end of the block.

##### Extracellular electrophysiological data acquisition in mice

Extracellular recordings were performed in V1 with microelectrode silicon probes. Full information about these recordings are detailed in the initial publication^37^. Briefly, on the day prior to extracellular recording sessions small craniotomies were made over V1 (relative to lambda: AP 0.0, ML ± 3.00 mm) and in a subset of animals also auditory cortex, posterior parietal cortex and medial prefrontal cortex. Only V1 data is included in this study. Prior to each session silicon microelectrode arrays with 32 or 64 channels were inserted into the cortex to span the layers of V1 and allowed to stabilize at least 15 minutes before starting data acquisition during task execution.

### Analysis of baseline pupil size

#### Mouse dataset – preprocessing of pupil data

Pupil size and position was extracted using DeepLabCut^76^. A neural network was trained on a representative excerpt of video frames with manually labeled pupil center and 6 marker points along the pupil outline. The remaining video data was subsequently automatically labeled. Full details about the extraction of pupil variables can be found in the original publication^37^. We converted the time series to units of modulation (percent signal change) around the median of the pupil time series from each block.

#### Human dataset – preprocessing of pupil data

Periods of blinks and saccades were detected using the manufacturer’s standard algorithms with default settings. The remaining data analyses were performed using custom-made Python software. We applied to each pupil recording (i) linear interpolation of values measured just before and after each identified blink (interpolation time window, from 150ms before until 150ms after blink), (ii) low-pass filtering below 10Hz), (iii) removal of pupil responses to blinks and to saccades, by first estimating these responses by means of deconvolution and then removing them from the pupil time series by means of multiple linear regression^77^, and (iv) conversion to units of modulation (percent signal change) around the median of the pupil time series from each block.

#### Quantification of baseline pupil size

In both datasets we calculated pre-trial pupil size measures as the mean pupil size during the 500 ms preceding stimulus (change) onset.

#### Session inclusion criteria

Experimental sessions were only included if pupillary recordings were performed and the average engaged-state probability during the session fell between 0.05 and 0.95. After applying these criteria, the dataset used for analyses linking behavior and pupil consisted of 10 mice, 74 sessions, and 26111 trials (**Figure 1**) and of 57 human participants (1 session per participant) and 41235 trials (**Figure 2**).

### Analysis of baseline V1 spiking activity

#### Preprocessing of extracellular electrophysiology data

Following the initial publication^37^, spike detection and sorting were performed using automatic clustering and posthoc manual curation using Klusta and the Phy GUI, respectively^78^. Only stable, high quality neurons were included for the final analysis when they had 1) an isolation distance higher than 10 (see^79^), 2) less than 0.1% of their spikes within the refractory period of 15ms, and 3) spiking activity in 90 out of 100 equally-sized time bins within an experimental session. To compute firing rates, spikes were aligned to the moment of stimulus change, binned in 10-ms bins and convolved with a causal half-Gaussian window with 50-ms standard deviation. All neurons with baseline (-500ms to 0 ms) or early evoked (0ms to 250ms) activity exceeding three standard deviations from the population mean were excluded from further analysis (**Figure S3A**).

#### Quantification of baseline V1 spiking activity

Individual neurons were classified as fast-spiking (FS) interneurons and putative pyramidal neurons based on peak-to-through delays^37^. Separately for each neuron in the dataset, we calculated baseline V1 spiking activity as the spike rate during the 500 ms preceding stimulus change (same as pupil size).

#### Session inclusion criteria

Experimental sessions were only included if simultaneous electrophysiological and pupillary recordings were performed, the average engaged-state probability during the session fell between 0.05 and 0.95, and at least three neurons (per neuron type) were recorded. After applying these criteria, the dataset used for analyses linking behavior, pupil and spiking activity (**Figure 3**) consisted of 6 mice, 17 sessions, 172 recorded putative pyramidal cells (4-21 per session), 95 recorded interneurons (3-11 per session), and 4698 trials.

### Behavioral state modeling using GLM-HMM

We used the Python package SSM^80^ to implement a multinomial GLM-HMM (for details, see ref^2,4^), and applied the same approach to the mouse and human datasets. In short, the dependent variable was choice, which could take one of three values on each trial: “auditory change”, “visual change”, or omission (mice) or “no”, “yes” or omission (humans). The GLM component modelled choices as a function of an intercept (capturing choice bias) and a stimulus input predictor (capturing sensitivity). For the mouse dataset, stimulus input on conflict trials (containing both auditory and visual stimulus changes) was determined based on the animals’ response and, when no response was provided, based on the parity of the trial number (even trials: auditory, odd trials: visual). The HMM component described a vector of initial probabilities of a fixed number of states that differed with respect to their GLM beta weights and a matrix of the average transition probabilities between these states. We fit the multinomial GLM-HMM to trials from individual subjects using the Expectation Maximization (EM) algorithm to maximize the log-posterior and obtain the optimized parameters.

Model selection for the number of states was performed using 5-fold cross-validation across sessions of an individual mouse and using 3-fold cross-validation across blocks of an individual human^2,4^. Models with up to seven states were fit to the concatenated trials of the training set, and we computed the log-likelihood of the held-out 20% of sessions using the fit parameters and a single run of the forward pass on the held-out sessions. Because the EM algorithm may lead to local maxima, for each choice of number of states, the EM algorithm was performed 10 times starting from random initial conditions. We report the log-likelihood of different models on held-out sessions in units of bits per trial^3^. The best number of states was chosen as the number of states for which adding one more state would not significantly increase the log-likelihood, which resulted in four states in mice (**Figure S1A**) and three states in human participants (**Figure S1B**). States were classified through visual inspection.

### Statistical analyses

#### GLM-HMM selection

To statistically evaluate improvements in model fit across different numbers of latent behavioral states, we compared cross-validated log-likelihoods of models using pairwise Wilcoxon rank tests (e.g., 1-state vs. 2-state models, etc.).

#### Linear mixed-effects models

To assess the relationship between trial-by-trial variations in baseline pupil size and engaged-state probability, while accounting for inter-subject variability, we fitted the following mixed-effects model on mouse and human data using the R package lme4^81^.

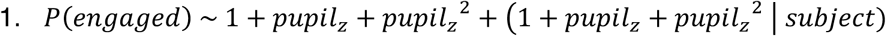

#### Nonlinear mediation analysis

A nonlinear mediation model was implemented R using Lavaan^82^, to assess whether the relationship between pupil-linked arousal and engaged-state probability was mediated by baseline spiking activity of V1 interneurons and putative pyramidal neurons. The mediation model was fit at the single-trial level with baseline V1 interneuron and pyramidal neuron activity as concurrent mediators. For each trial, pupil size (z-scored per session) was treated as predictor variable (𝑋), baseline firing rates (z-scored per neuron then averaged across neurons) of interneurons (𝑀*_i_*) and putative pyramidal neurons (𝑀*_p_*) as mediators, and engaged-state probability as the outcome (𝑌). All mediation paths were modelled with linear and quadratic terms to capture potential nonlinear relationships.

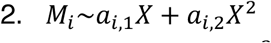

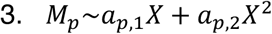

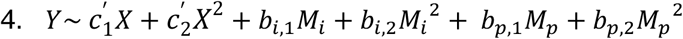

In nonlinear mediation models, the instantaneous indirect effect (𝐼𝐼𝐸) is defined as the product of the first partial derivative of M with respect to X and first partial derivative of output Y with respect to M, which can be expressed as a function of X^45^. Below, equations are shown for the IIE of interneurons (𝐼𝐸𝐸*_i_*), but the same equations apply to the IIE of pyramidal neurons.

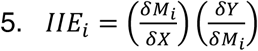

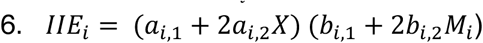

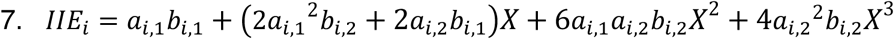

To establish the magnitude of the mediated effect, we calculated the mediated proportion of the total effect, with the total effect being the sum of both IIEs and the instantaneous direct effect (𝐼𝐷𝐸). The mediated proportion of the total effect (𝑃𝑀) was calculated per bootstrapped sample for each mediator across the pupil size range in which the mediation effects were significantly different between pyramidal neurons and interneurons (pupil ≥ 1.80). Note that this mediated proportion of the total effect can fall outside of the range 0-100% if the directions of the indirect and direct effects are in opposition. Below, the mediated proportion is defined for the proportion mediated by interneurons (𝑃𝑀*_i_*), but the same definitions apply to mediation by pyramidal neurons.

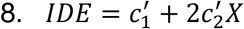

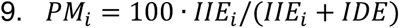

Statistical inference was performed via nonparametric bootstrapping (10,000 samples), performed at the level of single trials and not accounting for session differences. The two-sided p-value for each estimated parameter (β) was calculated as:

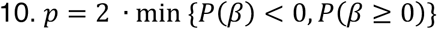

Statistical significance of the IIEs was assessed at each local pupil size value using the same p-value calculation described above. No cluster-based correction for multiple comparisons was applied, as the IIE is a continuous, deterministic function of a single nonlinear mediation model rather than a set of independent statistical tests. Values of the IIEs at different predictor levels are therefore not independent, but are jointly determined by the same model parameters.

## Data, Materials and Software availability

Data and analysis scripts will be made publicly available upon publication.

## Acknowledgments

### Author contributions

Conceptualization, SAN, SvG, JWdG; Investigation, SAN, MOL, ACS; Formal analysis, SAN, JWdG; Writing—original draft, SAN, JWdG; Writing—review and editing, MOL, ACS, SvG, UO, CP, PS; Supervision, PS, JWdG.

## Supplementary information

**Figure S1.**
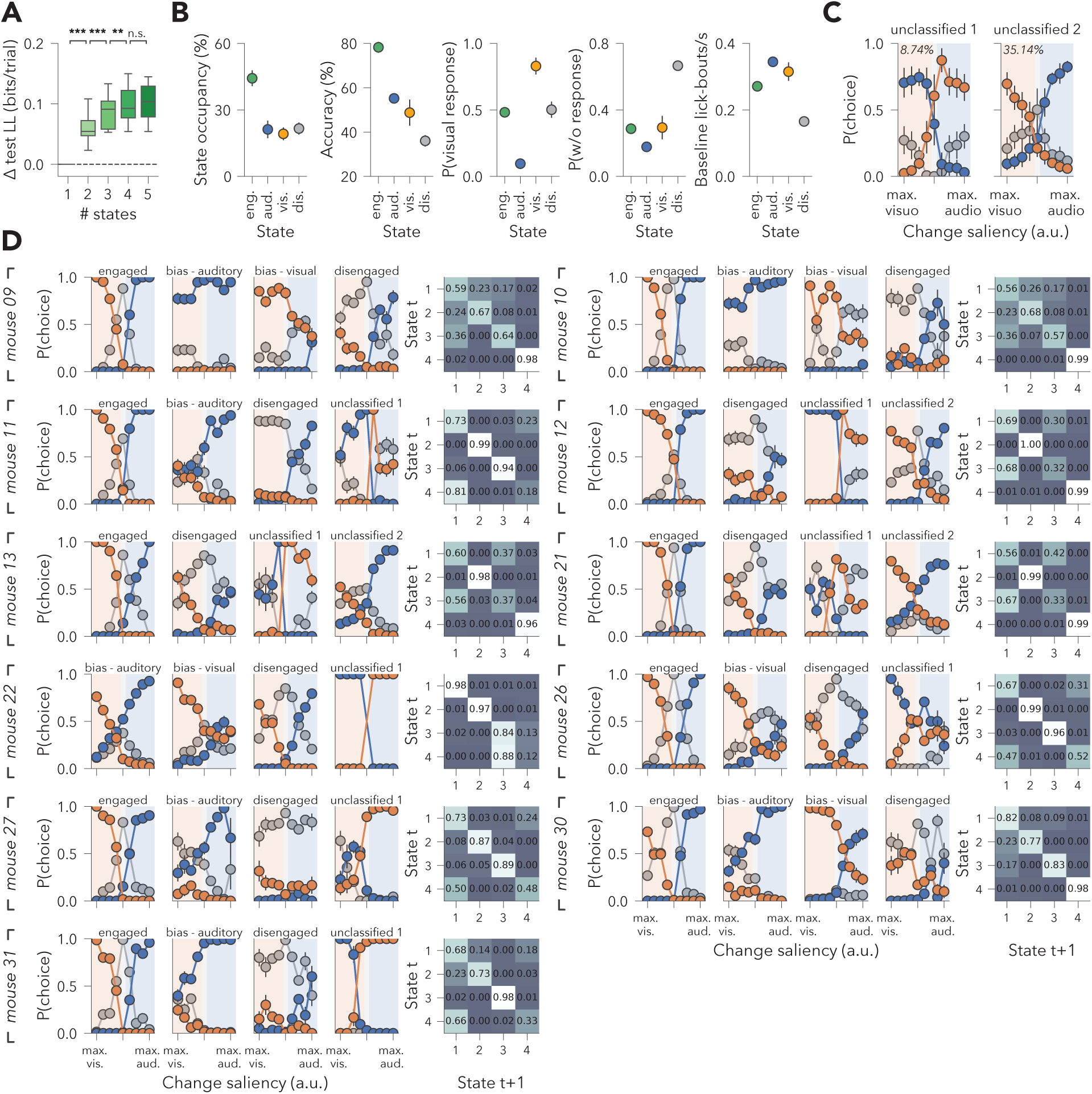
Model cross-validation, state descriptives, and individual-level model fits for mouse dataset. **(A)** Test-set log-likelihood values shown for models with one to five behavioral states, shown as change from a one-state model. Log-likelihoods increased with more states up to the four-state model, which was then selected for further analyses. **(B)** Average state occupancy, choice accuracy, visual response rates, behavioral lapses, and baseline lick-bout rates for the four classified behavioral states. Licks occurring within 0.5 s of one another were considered part of the same lick bout. **(C)** Two states were labelled as unclassified. Unclassified state 1 occurred in an average 8.74% of trials of eight mice and was characterized by inverted response-mappings or incoherent behavior. Unclassified state 2 occurred in an average 35.14% of trials of three mice and was characterized by unbiased, but low-accuracy decision-making. **(D)** Psychometric curves and transition matrices for the behavioral states of all animals. Error bars and shading indicate SEM across animals for panels B-C and SEM across trials for panel D. ***: p<.001, **: p<.01, n.s.: p>.05.

**Figure S2.**
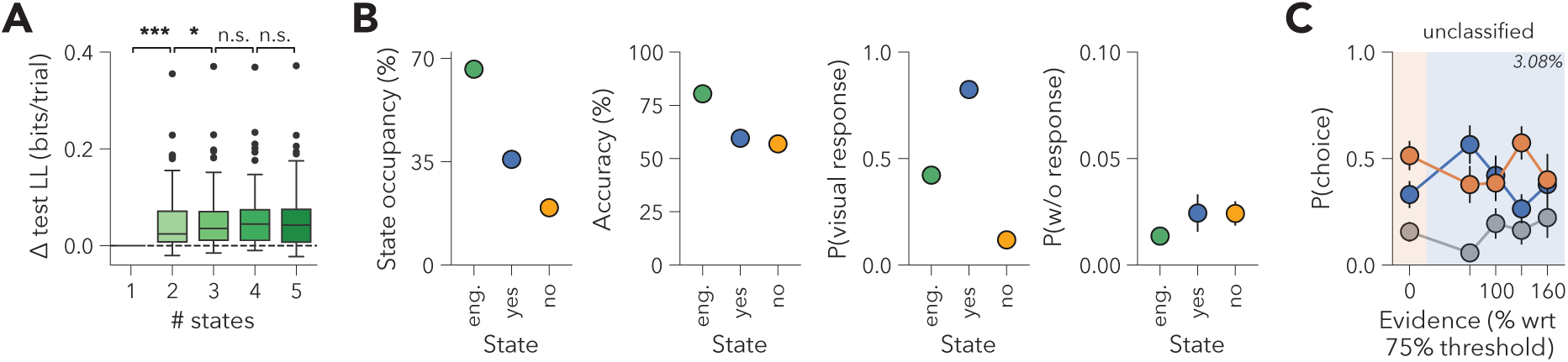
Model cross-validation and state descriptives for human dataset. **(A)** Test-set log-likelihood values shown for models with one to five behavioral states, shown as change from a one-state model. Log-likelihood values were compared between states using Wilcoxon signed-rank tests. Log-likelihoods increased with more states up to the three-state model, which was then selected for further analyses. **(B)** Average state occupancy, choice accuracy, visual response rates, and behavioral lapses for the three classified behavioral states. **(C)** One state was labelled as unclassified for the human dataset. This unclassified state occurred in an average 3.08% of trials of 36 humans and was not characterized by a coherent decision-making strategy. Error bars and shading indicate SEM across participants for panels B-C. ***: p<.001, *: p<.05, n.s.: p>.05.

**Figure S3.**
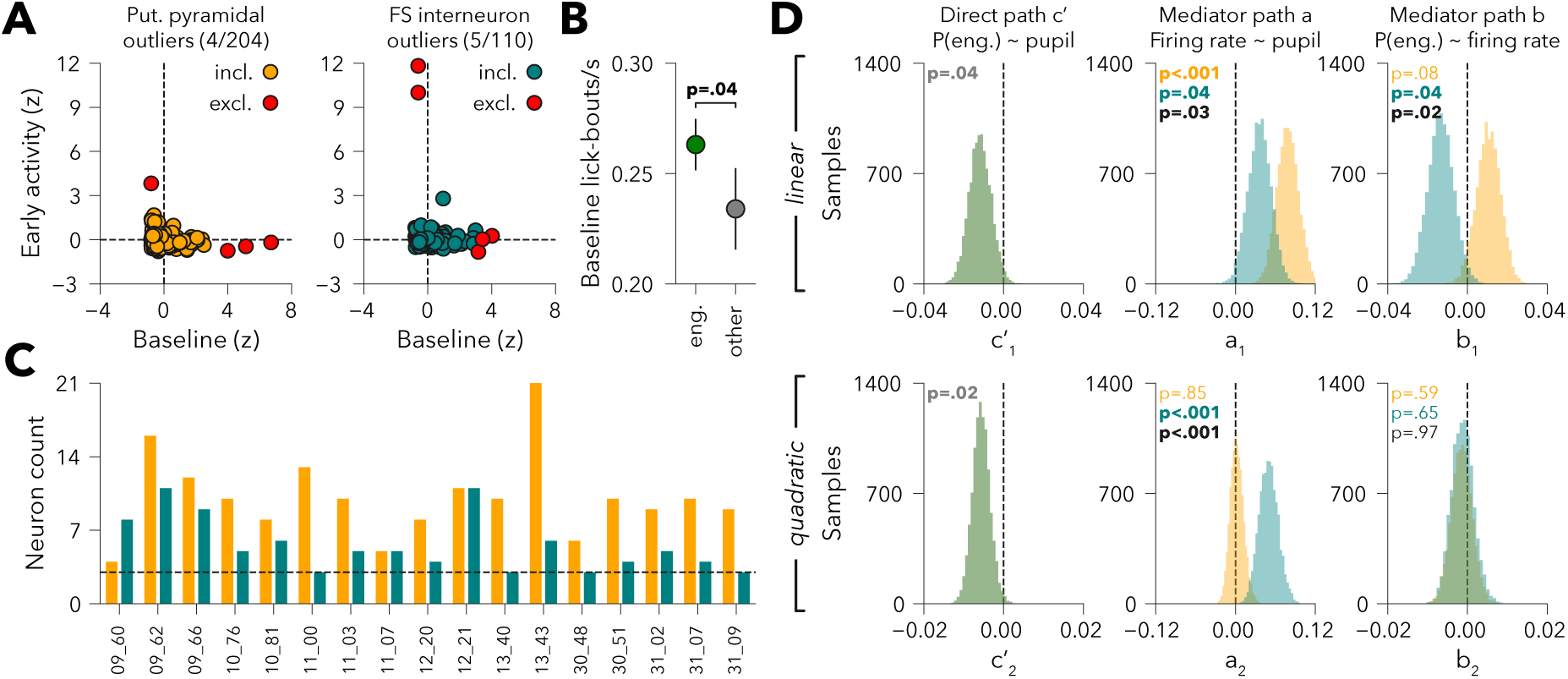
Data inclusion, licking behavior, and supplementary mediation results. **(A)** Neurons were excluded when baseline activity or early evoked activity exceeded three standard deviations from the full sample mean (across neuron types). For visualization purposes, included and excluded neurons are shown separately for putative pyramidal neurons (*left*; included in orange, excluded in red) and for interneurons (*right*; included in teal, excluded in red). The final dataset contains less neurons than reported in this figure, because several sessions with fewer than three neurons of each type were discarded after this preprocessing step. **(B)** Lick-bout rates were increased in the engaged state (in green) compared to all other non-engaged states (in grey) (paired-sample two-sided t-test on sessions; t(16)=2.20, p=.04). Error bars indicate SEM across sessions. **(C)** Neuron counts of included sessions. Sessions were only included when they contained at least three (dashed line) putative pyramidal neurons (in orange) and interneurons (in teal). **(D)** Bootstrapped (10,000 samples) linear (*top row*) and quadratic (*bottom row*) coefficients for paths c’, a, and b. Reported p-values correspond to two-sided tests against zero for each neuron type (putative pyramidal neurons in orange and interneurons in teal) and their difference (in black).

## References

1. Csikszentmihalyi, M. & Csikzentmihaly, M. Flow: The Psychology of Optimal Experience. vol. 1990 (Harper & Row New York, 1990).

2. Ashwood, Z. C. et al. Mice alternate between discrete strategies during perceptual decision-making. Nat Neurosci 25, 201–212 (2022).

3. Bolkan, S. S. et al. Opponent control of behavior by dorsomedial striatal pathways depends on task demands and internal state. Nat Neurosci 25, 345–357 (2022).

4. Hulsey, D., Zumwalt, K., Mazzucato, L., McCormick, D. A. & Jaramillo, S. Decision-making dynamics are predicted by arousal and uninstructed movements. Cell Reports 43, (2024).

5. Johnson, P. A., Nieuwenhuis, S., Mejías, J. & Urai, A. E. A dynamical systems model of arousal-driven behavioural state transitions. 2025.10.31.685593 Preprint at 10.1101/2025.10.31.685593 (2025).

6. Bandi, A. C. et al. Parietal cortex is causally required for state-dependent decisions. Cell Reports 44, 116672 (2025).

7. Weilnhammer, V., Chikermane, M. & Sterzer, P. Bistable perception alternates between internal and external modes of sensory processing. iScience 24, 102234 (2021).

8. Weilnhammer, V., Stuke, H., Standvoss, K. & Sterzer, P. Sensory processing in humans and mice fluctuates between external and internal modes. PLoS Biol 21, e3002410 (2023).

9. Weilnhammer, V. et al. N-methyl-d-aspartate receptor hypofunction causes recurrent and transient failures of perceptual inference. Brain 148, 1531–1539 (2025).

10. Schaub, A.-C., Eckert, A.-L., Nuiten, S. A., Weilnhammer, V. & Sterzer, P. Reduced weighting of short-term perceptual priors during auditory perceptual decision-making in psychosis-prone individuals. BMC Biol 23, 310 (2025).

11. Weilnhammer, V., Murai, Y. & Whitney, D. Dynamic predictive templates in perception. Curr Biol 34, 4301–4306.e2 (2024).

12. Aston-Jones, G. & Cohen, J. D. An Integrative Theory of Locus Coeruleus-Norepinephrine Function: Adaptive Gain and Optimal Performance. Annu. Rev. Neurosci. 28, 403–450 (2005).

13. Harris, K. D. & Thiele, A. Cortical state and attention. Nature Reviews Neuroscience 12, 509–523 (2011).

14. Lee, S.-H. & Dan, Y. Neuromodulation of brain states. Neuron 76, 209–222 (2012).

15. McCormick, D. A., Nestvogel, D. B. & He, B. J. Neuromodulation of Brain State and Behavior. Annu Rev Neurosci 43, 391–415 (2020).

16. Pfeffer, T. et al. Circuit mechanisms for the chemical modulation of cortex-wide network interactions and behavioral variability. Science Advances 7, eabf5620 (2021).

17. Froemke, R. C. Plasticity of Cortical Excitatory-Inhibitory Balance. Annu. Rev. Neurosci. 38, 195–219 (2015).

18. Joshi, S. & Gold, J. I. Pupil Size as a Window on Neural Substrates of Cognition. Trends in Cognitive Sciences 10.1016/j.tics.2020.03.005 (2020) doi:10.1016/j.tics.2020.03.005.

19. Reimer, J. et al. Pupil fluctuations track rapid changes in adrenergic and cholinergic activity in cortex. Nat Commun 7, 13289 (2016).

20. de Gee, J. W. et al. Dynamic modulation of decision biases by brainstem arousal systems. eLife 6, 309 (2017).

21. McGinley, M. J. et al. Waking State: Rapid Variations Modulate Neural and Behavioral Responses. Neuron 87, 1143–1161 (2015).

22. Beerendonk, L. et al. A disinhibitory circuit mechanism explains a general principle of peak performance during mid-level arousal. Proceedings of the National Academy of Sciences 121, e2312898121 (2024).

23. McGinley, M. J., David, S. V. & McCormick, D. A. Cortical Membrane Potential Signature of Optimal States for Sensory Signal Detection. Neuron 87, 179–192 (2015).

24. Papadopoulos, L. et al. Modulation of metastable ensemble dynamics explains the inverted-U relationship between tone discriminability and arousal in auditory cortex. Neuron 0, (2025).

25. Beerendonk, L. et al. Adaptive arousal regulation: Pharmacologically shifting the peak of the Yerkes–Dodson curve by catecholaminergic enhancement of arousal. Proceedings of the National Academy of Sciences 122, e2419733122 (2025).

26. Tong, C. et al. Norepinephrine-mediated arousal fluctuations drive inverted U-shaped functional connectivity dynamics. Nat Commun 10.1038/s41467-025-66436-x (2025) doi:10.1038/s41467-025-66436-x.

27. Gee, J. W. de et al. Strategic stabilization of arousal boosts sustained attention. Current Biology 34, 4114–4128.e6 (2024).

28. Yerkes, R. M. & Dodson, J. D. The relation of strength of stimulus to rapidity of habit-formation. Journal of Comparative Neurology and Psychology 18, 459–482 (1908).

29. Kim, J.-H. et al. Selectivity of Neuromodulatory Projections from the Basal Forebrain and Locus Ceruleus to Primary Sensory Cortices. J Neurosci 36, 5314–5327 (2016).

30. Poe, G. R. et al. Locus coeruleus: a new look at the blue spot. Nat Rev Neurosci 21, 644–659 (2020).

31. Pinto, L. et al. Fast modulation of visual perception by basal forebrain cholinergic neurons. Nat Neurosci 16, 1857–1863 (2013).

32. Polack, P.-O., Friedman, J. & Golshani, P. Cellular mechanisms of brain-state-dependent gain modulation in visual cortex. Nat Neurosci 16, 1331–1339 (2013).

33. Neske, G. T., Nestvogel, D., Steffan, P. J. & McCormick, D. A. Distinct Waking States for Strong Evoked Responses in Primary Visual Cortex and Optimal Visual Detection Performance. J Neurosci 39, 10044–10059 (2019).

34. Speed, A., Del Rosario, J., Burgess, C. P. & Haider, B. Cortical State Fluctuations across Layers of V1 during Visual Spatial Perception. Cell Rep 26, 2868–2874.e3 (2019).

35. Steinmetz, N. A., Zatka-Haas, P., Carandini, M. & Harris, K. D. Distributed coding of choice, action and engagement across the mouse brain. Nature 576, 266–273 (2019).

36. Jacobs, E. A. K., Steinmetz, N. A., Peters, A. J., Carandini, M. & Harris, K. D. Cortical State Fluctuations during Sensory Decision Making. Curr Biol 30, 4944–4955.e7 (2020).

37. Oude Lohuis, M. N., et al. Multisensory task demands temporally extend the causal requirement for visual cortex in perception. Nat Commun 13, 2864 (2022).

38. Oude Lohuis, M. N., Marchesi, P., Olcese, U. & Pennartz, C. M. A. Triple dissociation of visual, auditory and motor processing in mouse primary visual cortex. Nat Neurosci 27, 758–771 (2024).

39. Nuiten, S. A., et al. Phasic and tonic arousal distinctly shape human decision bias. Preprint at 10.21203/rs.3.rs-6479550/v1 (2025).

40. Nuiten, S. A., Gee, J. W. de, Zantvoord, J. B., Fahrenfort, J. J. & Gaal, S. van. Pharmacological Elevation of Catecholamine Levels Improves Perceptual Decisions, But Not Metacognitive Insight. eNeuro 11, (2024).

41. Yizhar, O. et al. Neocortical excitation/inhibition balance in information processing and social dysfunction. Nature 477, 171–178 (2011).

42. Lee, M., Mueller, A. & Moore, T. Differences in Noradrenaline Receptor Expression Across Different Neuronal Subtypes in Macaque Frontal Eye Field. Front Neuroanat 14, 574130 (2020).

43. Kaufman, K. J., Krall, R. F. & Williamson, R. S. Pupil-linked arousal differentially modulates cell-type-specific sensory processing. Preprint at 10.1101/2025.06.09.658645 (2025).

44. Del Rosario, J. et al. Lateral inhibition in V1 controls neural and perceptual contrast sensitivity. Nat Neurosci 28, 836–847 (2025).

45. Hayes, A. F. & Preacher, K. J. Quantifying and Testing Indirect Effects in Simple Mediation Models When the Constituent Paths Are Nonlinear. Multivariate Behavioral Research 45, 627–660 (2010).

46. de Gee, J. W. et al. Strategic stabilization of arousal boosts sustained attention. Current Biology 10.1016/j.cub.2024.07.070 (2024) doi:10.1016/j.cub.2024.07.070.

47. Cohen, J. D., McClure, S. M. & Yu, A. J. Should I stay or should I go? How the human brain manages the trade-off between exploitation and exploration. Philos Trans R Soc Lond B Biol Sci 362, 933–942 (2007).

48. Jepma, M. & Nieuwenhuis, S. Pupil diameter predicts changes in the exploration-exploitation trade-off: evidence for the adaptive gain theory. Journal of cognitive neuroscience 23, 1587–1596 (2011).

49. Arnsten, A. F. T. & Goldman-Rakic, P. S. Selective prefrontal cortical projections to the region of the locus coeruleus and raphe nuclei in the rhesus monkey. Brain Research 306, 9–18 (1984).

50. Joshi, S. & Gold, J. I. Context-dependent relationships between locus coeruleus firing patterns and coordinated neural activity in the anterior cingulate cortex. Elife 11, e63490 (2022).

51. Porrino, L. J. & Goldman-Rakic, P. S. Brainstem innervation of prefrontal and anterior cingulate cortex in the rhesus monkey revealed by retrograde transport of HRP. Journal of Comparative Neurology 205, 63–76 (1982).

52. Barkley, R. A. Behavioral inhibition, sustained attention, and executive functions: Constructing a unifying theory of ADHD. Psychological Bulletin 121, 65–94 (1997).

53. Zhao, S., Liu, Y. & Wei, K. Pupil-Linked Arousal Response Reveals Aberrant Attention Regulation among Children with Autism Spectrum Disorder. J. Neurosci. 42, 5427–5437 (2022).

54. de Lecea, L., Carter, M. E. & Adamantidis, A. Shining Light on Wakefulness and Arousal. Biological Psychiatry 71, 1046–1052 (2012).

55. Sander, C., Hensch, T., Wittekind, D. A., Böttger, D. & Hegerl, U. Assessment of Wakefulness and Brain Arousal Regulation in Psychiatric Research. NPS 72, 195–205 (2015).

56. Batista-Brito, R., Zagha, E., Ratliff, J. M. & Vinck, M. Modulation of cortical circuits by top-down processing and arousal state in health and disease. Current Opinion in Neurobiology 52, 172–181 (2018).

57. Murphy, P. R. et al. Individual differences in belief updating and phasic arousal are related to psychosis proneness. Commun Psychol 2, 1–14 (2024).

58. Kreis, I., Zhang, L., Moritz, S. & Pfuhl, G. Spared performance but increased uncertainty in schizophrenia: Evidence from a probabilistic decision-making task. Schizophrenia Research 243, 414–423 (2022).

59. Kreis, I. et al. Aberrant uncertainty processing is linked to psychotic-like experiences, autistic traits, and is reflected in pupil dilation during probabilistic learning. Cogn Affect Behav Neurosci 10.3758/s13415-023-01088-2 (2023) doi:10.3758/s13415-023-01088-2.

60. Kepecs, A., Uchida, N., Zariwala, H. A. & Mainen, Z. F. Neural correlates, computation and behavioural impact of decision confidence. Nature 455, 227–231 (2008).

61. Rushworth, M. F. S. & Behrens, T. E. J. Choice, uncertainty and value in prefrontal and cingulate cortex. Nat Neurosci 11, 389–397 (2008).

62. Joshi, S., Li, Y., Kalwani, R. M. & Gold, J. I. Relationships between Pupil Diameter and Neuronal Activity in the Locus Coeruleus, Colliculi, and Cingulate Cortex. Neuron 89, 221–234 (2016).

63. Murphy, P. R., O’Connell, R. G., O’Sullivan, M., Robertson, I. H. & Balsters, J. H. Pupil diameter covaries with BOLD activity in human locus coeruleus. Human brain mapping 35, 4140–4154 (2014).

64. Lloyd, B., de Voogd, L. D., Mäki-Marttunen, V. & Nieuwenhuis, S. Pupil size reflects activation of subcortical ascending arousal system nuclei during rest. Elife 12, e84822 (2023).

65. Varazzani, C., San-Galli, A., Gilardeau, S. & Bouret, S. Noradrenaline and Dopamine Neurons in the Reward/Effort Trade-Off: A Direct Electrophysiological Comparison in Behaving Monkeys. J. Neurosci. 35, 7866–7877 (2015).

66. Breton-Provencher, V. & Sur, M. Active control of arousal by a locus coeruleus GABAergic circuit. Nat Neurosci 22, 218–228 (2019).

67. Maheu, M., Donner, T. H. & Wiegert, J. S. Serotonergic and Noradrenergic Interactions in Pupil-linked Arousal. 2025.03.26.644382 Preprint at 10.1101/2025.03.26.644382 (2025).

68. Mridha, Z. et al. Graded recruitment of pupil-linked neuromodulation by parametric stimulation of the vagus nerve. Nat Commun 12, 1539 (2021).

69. Cazettes, F., Reato, D., Morais, J. P., Renart, A. & Mainen, Z. F. Phasic Activation of Dorsal Raphe Serotonergic Neurons Increases Pupil Size. Current Biology 31, 192–197.e4 (2021).

70. Brunton, B. W., Botvinick, M. M. & Brody, C. D. Rats and Humans Can Optimally Accumulate Evidence for Decision-Making. Science 340, 95–98 (2013).

71. de Gee, J. W. et al. Pupil-linked phasic arousal predicts a reduction of choice bias across species and decision domains. Elife 9, e54014 (2020).

72. Peters, E. R., Joseph, S. A. & Garety, P. A. Measurement of delusional ideation in the normal population: introducing the PDI (Peters et al. Delusions Inventory). Schizophr Bull 25, 553–576 (1999).

73. Klein, D. J., Depireux, D. A., Simon, J. Z. & Shamma, S. A. Robust Spectrotemporal Reverse Correlation for the Auditory System: Optimizing Stimulus Design. J Comput Neurosci 9, 85–111 (2000).

74. Peirce, J. et al. PsychoPy2: Experiments in behavior made easy. Behav Res 51, 195–203 (2019).

75. Kaernbach, C. Simple adaptive testing with the weighted up-down method. Perception & Psychophysics 49, 227–229 (1991).

76. Mathis, A. et al. DeepLabCut: markerless pose estimation of user-defined body parts with deep learning. Nat Neurosci 21, 1281–1289 (2018).

77. Knapen, T. et al. Cognitive and Ocular Factors Jointly Determine Pupil Responses under Equiluminance. PLOS ONE 11, e0155574 (2016).

78. Rossant, C. et al. Spike sorting for large, dense electrode arrays. Nat Neurosci 19, 634–641 (2016).

79. Schmitzer-Torbert, N., Jackson, J., Henze, D., Harris, K. & Redish, A. D. Quantitative measures of cluster quality for use in extracellular recordings. Neuroscience 131, 1–11 (2005).

80. Linderman, S., Antin, B., Zoltowski, D. & Glaser, J. Bayesian Learning and Inference for State Space Models. (2020).

81. Bates, D., Mächler, M., Bolker, B. & Walker, S. Fitting Linear Mixed-Effects Models Using lme4. Journal of Statistical Software 67, 1–48 (2015).

82. Rosseel, Y. lavaan: An R Package for Structural Equation Modeling | Journal of Statistical Software. Journal of Statistical Software 48, 1–36 (2012).

